# *Escherichia coli* isolated from commercial layer hens in Australia remain susceptible to critically important antimicrobials

**DOI:** 10.1101/2023.02.03.526949

**Authors:** Rebecca Abraham, Hui San Allison, Terence Lee, Anthony Pavic, Raymond Chia, Kylie Hewson, Zheng Z Lee, David J Hampson, David Jordan, Sam Abraham

## Abstract

Controlling the use of the most critically important antimicrobials (CIAs) in food animals has been identified as one of the key measures required to curb the transmission of antimicrobial resistant bacteria from animals to humans. Expanding the evidence demonstrating the effectiveness of restricting CIA usage for preventing the emergence of resistance to key drugs amongst commensal organisms in animal production would do much to strengthen international efforts to control antimicrobial resistance (AMR). As Australia has strict controls on antimicrobial use in layer hens, and internationally comparatively low levels of poultry disease due to strict national biosecurity measures, we investigated whether these circumstances have resulted in curtailing development of critical forms of AMR. The work comprised a cross-sectional national survey of 62 commercial layer farms with each assessed for AMR in *Escherichia coli* isolates recovered from faeces. Minimum inhibitory concentration analysis using a panel of 13 antimicrobials was performed on 296 isolates with those exhibiting phenotypic resistance to fluoroquinolones (a CIA) or multi-class resistance (MCR) subjected to whole genome sequencing. Overall, 52.0% of the isolates were susceptible to all antimicrobials tested, and all isolates were susceptible to ceftiofur, chloramphenicol and colistin. Resistance was observed to ampicillin (16.2%), cefoxitin (1.4%), ciprofloxacin (2.7%), florfenicol (2.4%), gentamicin (1.0%), streptomycin (4.7%), tetracycline (37.8%) and trimethoprim/sulfamethoxazole (10.5%). Multi-class resistance was observed in 23 isolates (7.7%), with one isolate (ST746) exhibiting resistance to five antimicrobial classes. Whole genome sequencing found that ciprofloxacin-resistant (fluoroquinolone) isolates were devoid of both known chromosomal mutations in the quinolone resistance determinant regions and plasmid-mediated quinolone resistance genes (*qnr*) - other than in one isolate (ST155) which carried the *qnrS* gene. Two MCR *E. coli* isolates with ciprofloxacin-resistance were found to be carrying known resistance genes including *aadA1, dfrA1, strA, strB, sul1, sul2, tet(A), bla*_TEM-1B_*, qnrS1* and *tet(A*). Overall, this study found that *E. coli* from layer hens in Australia have low rates of AMR, likely due to strict control on antimicrobial usage achieved by the sum of regulation and voluntary measures.

## Introduction

Resistance amongst bacteria to high priority, critically important antimicrobials (CIAs) – such as extended-spectrum cephalosporins (ESCs), fluoroquinolones (FQ), carbapenems, and colistin – threatens the therapeutic options for treatment of severe infections in humans (1). Emergence of CIA-resistant bacteria in food-producing animals further exacerbates this risk, either due to bacteria (commensals or pathogens) or genetic determinants being transferred via food, or via environmental pathways to infiltrate human microflora. Globally, there have been increasing reports of CIA-resistant indicator bacteria such as *Escherichia coli* isolated from food-producing animals. In 2020, the Danish Integrated Antimicrobial Resistance Monitoring and Research Programme (DANMAP) reported a 16% prevalence of FQ-resistant *E. coli* in broiler chickens (2) while a recent French study reported a 16.5% prevalence of colistin-resistant *E. coli* in veal calves (3). Moreover, in 2019 the American National Antimicrobial Resistance Monitoring System (NARMS) reported a 3.5% prevalence of FQ-resistant *E. coli* in pigs which was the highest reported prevalence to date in the United States (4).

One exception to the trend for increasingly prevalent CIA colonisation of food-animals is the Australian livestock sector (5–10). This favourable antimicrobial resistance (AMR) status has been attributed to Australia’s isolated geographic location, longstanding regulatory constraints on the use of CIAs (such as ESCs, FQs and colistin) in food animals (11) combined with strict quarantine conditions at the national border (12). Some sectors also achieve avoidance of antimicrobials by virtue of both the limited impact of bacterial disease and the constrained availability of antimicrobials as part of the regulatory process for eliminating chemical residues from animal products. An example of the latter is the commercial layer hen industry where for much of a hen’s existence, they are continuously producing eggs for human consumption and therefore must be minimally exposed to medications, including antimicrobials, that might accumulate residues in eggs.

Eggs from the domestic fowl are one of the most heavily consumed animal products worldwide. In Australia, there has been strong growth in consumer demand for eggs over the past decade. In 2021 it was estimated that on-average 246 eggs were consumed per person (13). This increasing level of egg consumption, has been linked to increases in foodborne illness outbreaks, often associated with sub-optimal handling of eggs in food establishments (14). Further, the use of poultry manures as fertiliser on horticultural crops can potentially produce other avenues for foodborne illness in humans (15). To improve the clarity of the AMR risks from the egg industry to the community, surveillance of the AMR status of layer flocks is needed. The most recent survey investigated AMR presence in *Salmonella* (16), considered in the context of previous data (17), revealed overall low rates of resistance to all tested antimicrobials. However, for the purpose of comparison, and for establishing a baseline description of AMR in the layer hen sector, a comprehensive evaluation of the AMR status of *E. coli* in layer hens is needed. In this work we hypothesised that *E. coli* from Australian layer hens would display low rates of AMR due to longstanding restrictions on antimicrobial use that prevail in the livestock sector, low levels of AMR detected in previous surveys of other bacteria and because of the special constraints applying to antimicrobial use in layer hens. The hypothesis was explored with a cross-sectional survey on a national scale which aimed to define the AMR carriage among commensal *E. coli* isolates from commercial layer hens. Genomic characterisation of isolates expressing resistance to CIA and expressing resistance to multiple classes (MCR) were undertaken to support inferences on the epidemiological significance of findings.

## Materials and Methods

### Sample Collection and Approach

Cloacal swabs (n=296) were collected from healthy commercial table egg laying chickens on 62 enrolled farms between August 2019 and January 2020. Five swabs were collected from each production unit, defined by management system (e.g. caged, free-range egg production systems). A commercial entity was eligible in this survey if they had at least one commercial egg production unit. This work was approved by Murdoch University Ethics committee (Permit number: Cadaver 944).

### *E. coli* isolation

Swabs were vortexed in buffered peptone water followed by direct streaking onto *E. coli* chromogenic agar (Chromagar ECC, Edwards Australia). The agar plates were incubated at 37 °C for 18 hours and then one colony was selected and sub-cultured onto Coli ID for purity. *E. coli* isolation was confirmed using an indole test followed by MALDI-TOF MS (Microflex, Bruker, MA, USA) prior to antimicrobial susceptibility testing.

### Antimicrobial Susceptibility Testing

The susceptibility of the isolates to antimicrobials was determined by broth microdilution according to the Clinical and Laboratory Standards Institute (CLSI) ISO 20776 standards (18). Drug panels were prepared in-house using a customised Freedom EVO genomics platform (19). Inocula were prepared manually and dispensed using an auto-inoculator (Sensititre). After incubation at 37 °C for 18 hours, images of the assay plates were taken, and the results recorded using the Sensititre Vizion system. *E. coli* ATCC 25922 was used as the control strain. The antimicrobials tested were amoxicillin/clavulanic acid, ampicillin, cefoxitin, ceftiofur, ceftriaxone, chloramphenicol, ciprofloxacin, florfenicol, gentamicin, colistin, streptomycin, tetracycline and trimethoprim/sulfamethoxazole.

Interpretation of susceptibility was based on The European Committee on Antimicrobial Susceptibility Testing (EUCAST) epidemiological cut-off (ECOFF) breakpoints which categorise isolates as wild type or non-wild (NWT) type (20) and differs from CLSI clinical breakpoints that determines whether isolates are clinically resistant to the antimicrobial used for clinical treatments (21). Multi-class resistance (MCR) is classified by ECOFF (NWT) for isolates exhibiting NWT phenotype to three or more antimicrobial classes (≥ 3 classes of antimicrobials). Clinical breakpoints are used to classify resistance and MCR in the absence of ECOFF break points which includes amoxicillin-clavulanate and ceftriaxone (21).

### Whole genome sequencing

Whole genomic analysis was undertaken for any *E. coli* isolates exhibiting resistance to CIAs. DNA extraction was performed on the isolates using the MagMAX Multi-sample DNA extraction kit (ThermoFisher Scientific) as per the manufacturer’s instructions. DNA library preparation was performed using a Celero™ DNA-Seq Library Preparation Kit Preparation kit (NuGen, Tecan) and then was sequenced on the Illumina Nextseq platform (22).

### Bioinformatics analysis

Sequencing files were assembled using SPAdes denovo assembler (v3.14.0) (23). Antimicrobial resistance genes (ARGs) were identified using ABRicate (v0.8.7) with default settings. The sequence types (STs) of the sequenced isolates were established by *in silico* multilocus sequence typing (MLST) as previously described (22).

## Results

### Phenotypic antimicrobial resistance

In total, 296 *E. coli* isolates were recovered from birds on 62 farms. Majority of the isolates were susceptible to all antimicrobials tested (*n* = 154, 52.0%), and all isolates were susceptible to ceftiofur, ceftriaxone, chloramphenicol and colistin (**Figure 1, Table 1**). Resistance was observed for ampicillin (16.2%), amoxicillin-clavulanate (2.7%), cefoxitin (1.4%), ciprofloxacin (2.7%), florfenicol (2.4%), gentamicin (1.0%), streptomycin (4.7%), tetracycline (37.8%) and trimethoprim/sulfamethoxazole (10.5%) (**Table 1**). Of the eight isolates classified as NWT for ciprofloxacin (fluoroquinolone, CIA) (**Table 1**), none were also above the clinical breakpoint of 1 mg/L.

**Figure 1.**
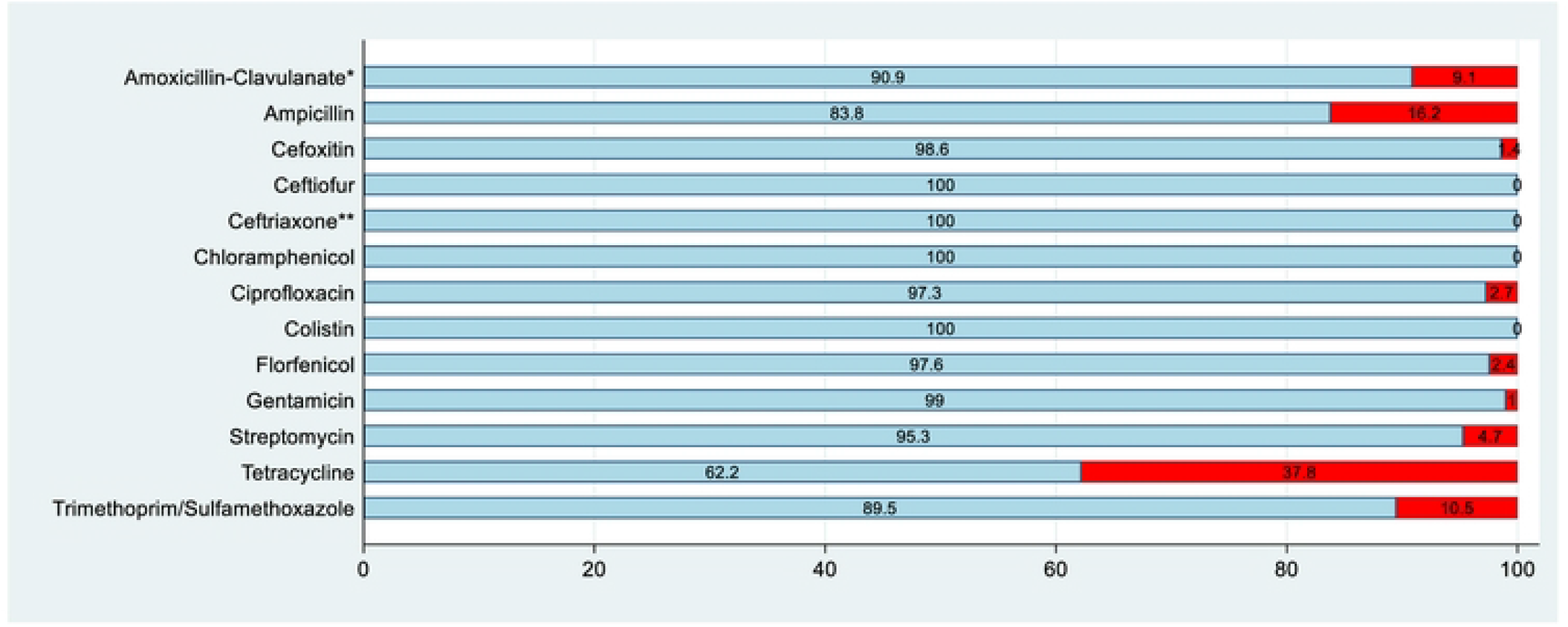
Antimicrobial resistance patterns for *E. coli* (*n* = 296) based on EUCAST epidemiological cutoff (ECOFF) breakpoints. The percentage of susceptible isolates is marked in blue and the percentage of resistant isolates is marked in red, unless otherwise indicated by footnotes (Percentage results were rounded to one decimal place). * Data for this antimicrobial was based on CLSI clinical breakpoints due to a lack of ECOFF breakpoint. ** Data for this antimicrobial was based on CLSI clinical breakpoints due to the ECOFF breakpoint being below the dilution range.

**Table 1.**
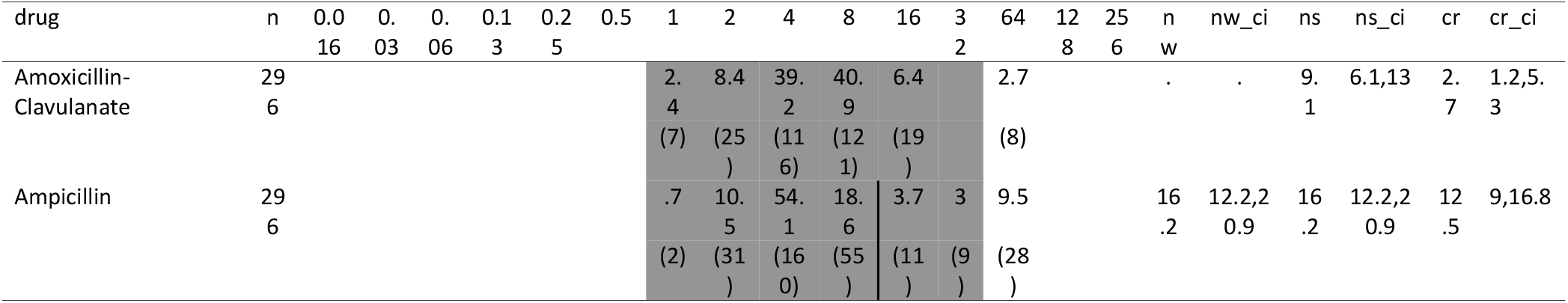
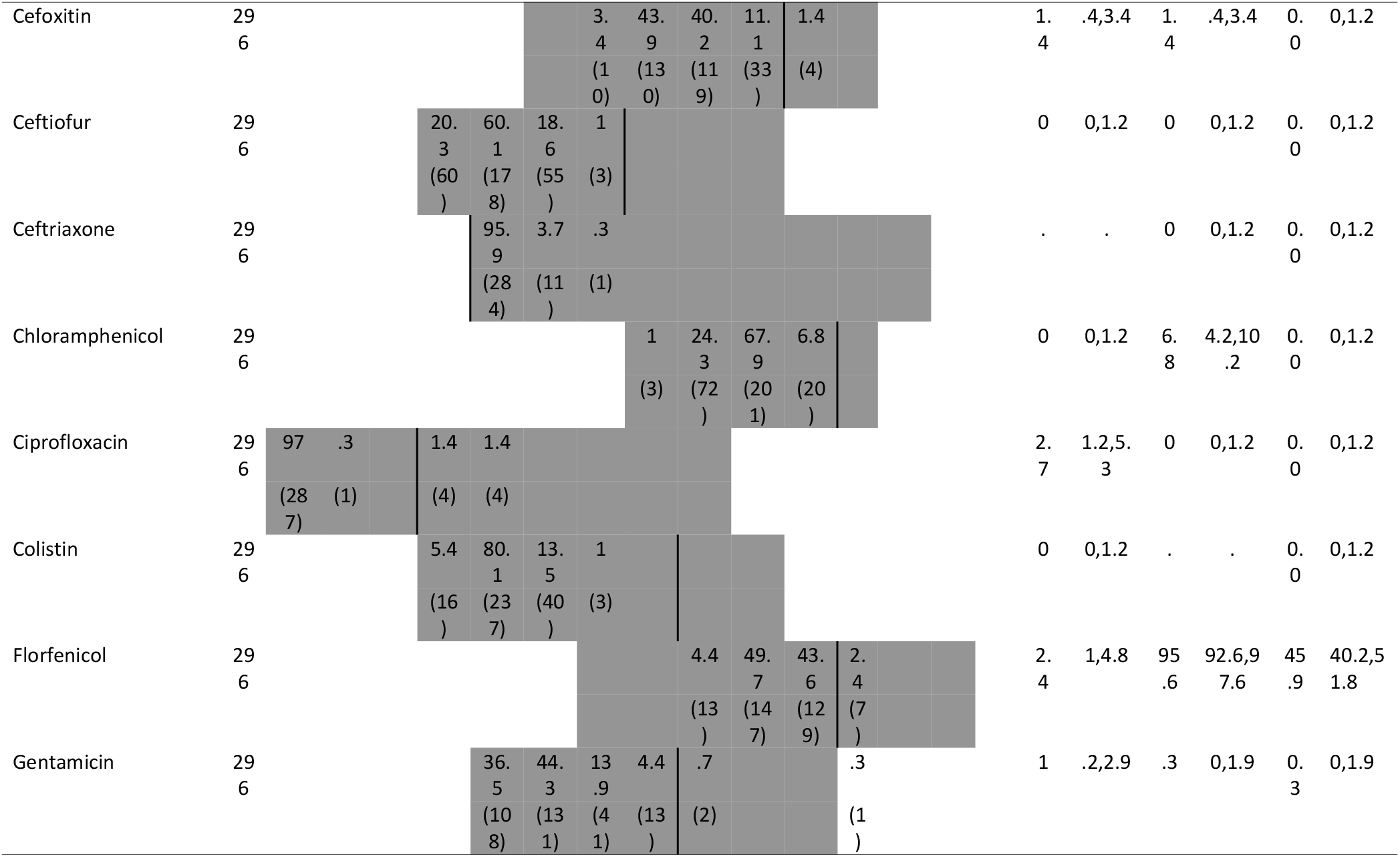
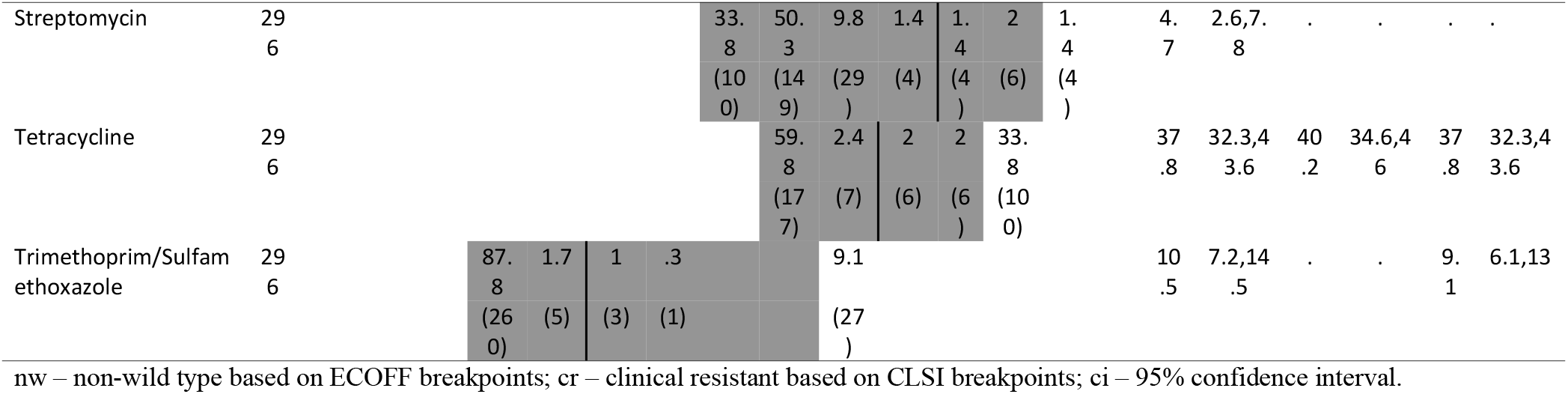
Distribution of minimum inhibitory concentrations (mg/L) for *Escherichia coli* isolates (*n* = 296) from faeces of Australian layer chickens. Percentage of resistance based on ECOFF breakpoints (non-wild type) and clinical breakpoints (clinical resistance) are presented on the right side (Percentage results were rounded to one decimal place). Vertical bars indicate position of ECOFF breakpoint if available.

### *Multi-class resistance profiles for* E. coli

Twenty-three (7.7%) *E. coli* isolates were identified as being MCR, with seven MCR profiles identified (**Table 2**). The most common MCR profile was beta-lactamases, folate pathway inhibitors and tetracyclines (*n* = 12, 4.1%). One isolate (0.3%) demonstrated resistance to four classes of antimicrobials (aminoglycosides, beta-lactams, folate pathway inhibitors and tetracyclines) while another isolate (0.3%) demonstrated resistance to five antimicrobial classes (aminoglycosides, first-generation cephalosporins, beta-lactams, folate pathway inhibitors and tetracyclines).

**Table 2.** Class-based antimicrobial susceptibility profiles of *E. coli* isolates (*n* = 296) obtained from Australian layer chickens.

| phenotype | n | pct |
| --- | --- | --- |
| 0: nil | 154 | 52.0 |
| 1: ami | 2 | 0.7 |
| 1: bla | 11 | 3.7 |
| 1: c1g | 3 | 1.0 |
| 1: fpi | 3 | 1.0 |
| 1: phe | 4 | 1.4 |
| 1: qui | 6 | 2.0 |
| 1: tet | 52 | 17.6 |
| 2: ami_tet | 6 | 2.0 |
| 2: bla_phe | 1 | 0.3 |
| 2: bla_tet | 20 | 6.8 |
| 2: fpi_tet | 11 | 3.7 |
| 3: ami_bla_tet | 3 | 1.0 |
| 3: ami_fpi_tet | 3 | 1.0 |
| 3: bla_fpi_tet | 12 | 4.1 |
| 3: bla_phe_tet | 2 | 0.7 |
| 3: bla_qui_tet | 1 | 0.3 |
| 4: ami_bla_fpi_tet | 1 | 0.3 |
| 5: ami_c1g_fpi_qui_tet | 1 | 0.3 |

Phenotype based on ECOFF breakpoints. ami= aminoglycosides, bla= beta lactams, c1g=1st generation cephalosporins, fpi= folate pathway inhibitors, , qui = quinolones, tet= tetracyclines

### *Genomic characterisation of critically important antimicrobial resistant* E. coli

Whole genome sequencing was undertaken on all eight *E. coli* isolates exhibiting phenotypic resistance to fluoroquinolones (CIA), two of which showed MCR phenotypes (**Table 3**). QRDR mutations or AMR genes for fluoroquinolone resistance were only detected in one isolate that carried *qnrS1*, a known plasmid-mediated quinolone-resistance gene (PMQR) (24). The two MCR *E. coli* isolates harbouring phenotypic quinolone resistance belonged to two different sequence types (ST746 and ST155) and carried AMR genes to first-line antimicrobials such as tetracycline and aminoglycosides. Not all phenotypically detected resistance was confirmed by the detection of corresponding AMR genes or mutations, a situation that has previously been described (25, 26).

**Table 3:** Phenotype and genotype data for quinolone-resistant *E. coli* isolates.

| MLST | Phenotype | Genotype | Isolates (n) |
| --- | --- | --- | --- |
| ST746 | c1g, qui, ami, tet, fpi | *aadA1, dfrA1, strA, strB, sul1, sul2, tet(A)* | 1 |
| ST155 | qui | *lnu(C)* | 1 |
| ST3714 | qui | *sul1* | 1 |
| ST155 | bla, qui, tet | *blaTEM-1B, qnrS1, tet(A)* | 1 |
| ST155 | qui | - | 2 |
| ST355 | qui | - | 2 |

Phenotype based on ECOFF breakpoints. ami= aminoglycosides, bla= beta lactams, c1g=1st generation cephalosporins, fpi= folate pathway inhibitors, qui = quinolones, tet= tetracyclines

## Discussion

This study demonstrated that there is an overall low level of AMR in *E. coli* isolated from Australian layer hens compared to other countries (27–30), with more than half of all isolates (52.0%) being susceptible to all tested antimicrobials, and with low levels (2.7%) or absence of resistance to CIAs (FQs, ESCs, colistin). MCR phenotypes were observed only among a small number of *E. coli* isolates (7.7%), with one isolate exhibiting resistance to five antimicrobial classes (aminoglycosides, first-generation cephalosporins, folate pathway inhibitors, quinolones and tetracyclines). Where phenotypic results suggested resistance, this was not supported by the presence of known resistance genes, except in a single isolate that harboured a known PMQR gene. Overall, the findings are consistent with other recent Australian studies demonstrating overall low levels of resistance to CIAs among *E. coli* isolated from Australian livestock (9, 17, 31).

As layer hens are often excluded from national AMR surveillance systems (unlike broilers), there has been relative paucity of drug susceptibility data for *E. coli* from this class of livestock. However, comparison of findings from the current study with recent international AMR studies in layer hens shows an overall lower level of *E. coli* resistance among Australian layer hens compared to other countries. For example, ampicillin-resistant *E. coli* was only detected among 16.2% of isolates from this current study compared to 83% and 49.5% for China (30) and Tanzania (28) respectively, while tetracycline-resistant *E. coli* was found among 37.8% of isolates from this current study compared to 87.3% and 53.6% for China (30) and Austria (27) respectively. This also extends to CIA-resistant *E. coli* where the current study demonstrated low levels or absence of CIA-resistant *E. coli* among Australian layer hens compared to China (30), Austria (27) and Tanzania (28). Examples include low levels of cefoxitin-resistance (1.4%) compared to Austria (56%), low levels of ciprofloxacin-resistance (2.7%) compared to China (45.8%) and Tanzania (33.8%), and the absence of colistin resistance compared to China (4.9%) and Austria (73.7%).

Some resistance genes encountered in this study were directed against antimicrobials that are not approved for use in Australian layer hens (eg. fluroquinolones). This phenomenon reflects the labile nature of resistance in *E. coli* populations arising from an acquisition of resistance by gene-transfer and co-selection, adaptations allowing survival in the environment between host colonisation events, and ability to colonise multiple host species of which some are highly mobile and can spread resistance between populations. The findings in this respect are consistent with some other recent studies demonstrating low levels of AMR to high or medium importance antimicrobials among *E. coli* isolated from Australian cattle and meat chickens (8, 32). In addition, there is increasing empirical data revealing cross-species transfer of resistant *E. coli* from humans or other animals such as pigs (33), wild birds (34, 35) and poultry (36) regardless of which antimicrobials the recipient species are exposed to (22). It is possible that these transmission pathways have resulted in layer hens being exposed to and colonised with resistant bacteria. Another finding that requires explanation is the lack of both chromosomal QRDR mutations and PMQR (*qnr*) genes among all but one of the ciprofloxacin-resistant isolates and the lack of resistance based on clinical breakpoints indicate either MIC drift or the insensitivity of ECOFF breakpoints to distinguish wild type versus non-wild type phenotypes, as previously reported. Future investigations should consider the prevalence of resistance to antimicrobials of animal health significance to ensure One Health approaches to AMR are implemented to avoid unintended animal health consequences from only focusing on antimicrobials of importance to human health, and to continue work on minimising flock disease and adopting alternative therapies that do not require antimicrobial use.

Overall, this study supports similar findings from recent studies of Australian cattle, pigs and meat chickens (8–10, 16, 32, 37). The results indicate that significant resistance to antimicrobials of high importance were absent in commensal *E. coli* isolated from Australian layer hens. Where in some instances phenotypic results suggested resistance to CIA’s, this was not supported by the concurrent presence of known resistance genes in those same isolates, except in a single isolate that harboured a known quinolone resistance gene. This reflects the success of decades of stringent regulatory control on antimicrobial use, biosecurity measures, and infection prevention practises within the Australian egg industry. Altogether, the data here highlights the importance of understanding the transmission of resistant bacteria amongst all hosts (including humans) and to use biosecurity barriers to avoid even minor uses of antimicrobials that could lead to amplification and dispersal of unwanted forms of resistance.

## Acknowledgements

This project was funded by the Australian Department of Agriculture, Fisheries and Forestry, and Australian Eggs.

